# Ancient Truncated FtsZ Paralogs Likely Tune Cell Division in Hyphomicrobiales

**DOI:** 10.1101/2025.09.29.679267

**Authors:** Brody Aubry, Amelia Randich, Bailey Hudson, Elizabet Horton, Pamela J.B. Brown

## Abstract

Bacterial cell division is a well conserved, tightly regulated process that allows the separation of two viable daughter cells. In most bacteria, the proteins that drive division, termed the divisome, are recruited to mid-cell by FtsZ after it polymerizes to form the Z-ring. Interestingly, FtsZ has undergone several independent duplication events across the bacterial kingdom. We identified FtsZ GTPase protein sequences across alphaproteobacterial genomes, from representative genera for each family, and observed numerous *ftsZ* duplications in the order Hyphomicrobiales. Hidden Markov Modeling (HMM) supported the maintenance of two distinct lineages of FtsZ GTPase duplications among three families. The *Nitrobacteraceae* duplication, occurring in only the genus *Bradyrhizobium*, exhibits a different substitution pattern from that shared by the *Phyllobacteraceae* and *Rhizobiaceae* families. Within the *Rhizobiaceae* lineage, *Agrobacterium tumefaciens* contains three paralogs of FtsZ including the essential FtsZ_AT,_ and paralogs FtsZ_1_, and FtsZ_3_. In *A. tumefaciens,* we show that FtsZ_1_, but not FtsZ_3_, inhibits cell division when overexpressed. A hyperactive allele of *ftsW* partially protected against overexpression of *ftsZ_1_* suggesting that FtsZ_1_ may inhibit proper regulation of septal peptidoglycan biosynthesis during cell division. Overall, these observations suggest that maintenance of FtsZ paralogs in some bacteria which may fine tune the division process.

**Article Summary:** Gene duplication drives evolutionary innovation by providing raw genetic material and allowing microbes to acquire novel functions. Here, we explore the duplication of a gene encoding the essential cell division protein FtsZ in the Hyphomicrobiales and find that the duplication is primarily conserved in genera that interact with plant hosts. In *A. tumefaciens* FtsZ_1_ is not essential for cell division; however, overexpression of FtsZ_1_ inhibited cell division suggesting it may have a regulatory role during this essential process. This observation highlights the key role for gene duplication in the modulation of complex processes in bacteria.

## Introduction

Nearly all bacteria depend on the ancient cytoskeletal scaffold protein FtsZ to enable cell division and produce the next generation of cells (Megrian et al. 2022). This protein forms the “Z-ring,” and recruits a large complex of divisome proteins necessary to orchestrate cell division (Radler & Loose, 2024). While most bacteria have a single copy of *ftsZ*, at least five independent duplications have been observed across the bacterial kingdom (Vaughan et al. 2004; Margolin, 2005; Howell et al. 2019). Interestingly, these duplication events are not confined to one class of bacteria and the *ftsZ* paralogs are often truncated. Despite the distribution and frequency of *ftsZ* duplications, little is known about why these truncated *ftsZ* genes are maintained in genomes and if they encode novel functions. Here, we sought to expand our knowledge about the function of FtsZ paralogs within the Alphaproteobacterial class of bacteria.

Our understanding of FtsZ and its role in cell division is built on decades of work studying full length, “canonical” *ftsZ* that is encoded in the <u>d</u>ivision and <u>c</u>ell <u>w</u>all (*dcw*) operon alongside other essential components for division (Megrian et al. 2022). The FtsZ ring forms at mid-cell by self-oligomerization leading to the formation of dynamic filaments that are anchored to the inner membrane. The Z-ring forms a scaffold for the recruitment of division proteins, forming the mature divisome. Collectively, the divisome components complete four major steps of bacterial cell division: formation/anchoring of the Z-ring, recruitment of divisome proteins, activation of septal peptidoglycan synthases (FtsW/FtsI), and separation of daughter cells (Santiago-Collazo et al. 2024). Full length canonical *ftsZ* sequences possess four structural domains: N-Terminal Peptide (NTP) region, GTPase domain, C-Terminal Linker (CTL), and C-Terminal Domain (CTD) (Figure 1, grey box). The GTPase domain drives the dynamic self- oligomerization mediated by the binding and hydrolysis of GTP (de Boer et al. 1992; Mukherjee & Lutkenhaus, 1994; Erickson et al. 1996), the CTD is involved in anchoring (Ma et al. 1996; Addinall et al. 1996; Wang et al. 1997), and the CTL has roles in protein interactions and divisome regulation (Buske & Levin, 2013; Huecas et al. 2017; Sundararajan & Goley, 2017; Barrows et al. 2020). Lastly, the NTP, although more variable and less understood, has been shown to interact with regulating partners (Huang et al. 1996; Tan et al. 2010; Corrales-Guerrero, 2018).

**Figure 1.**
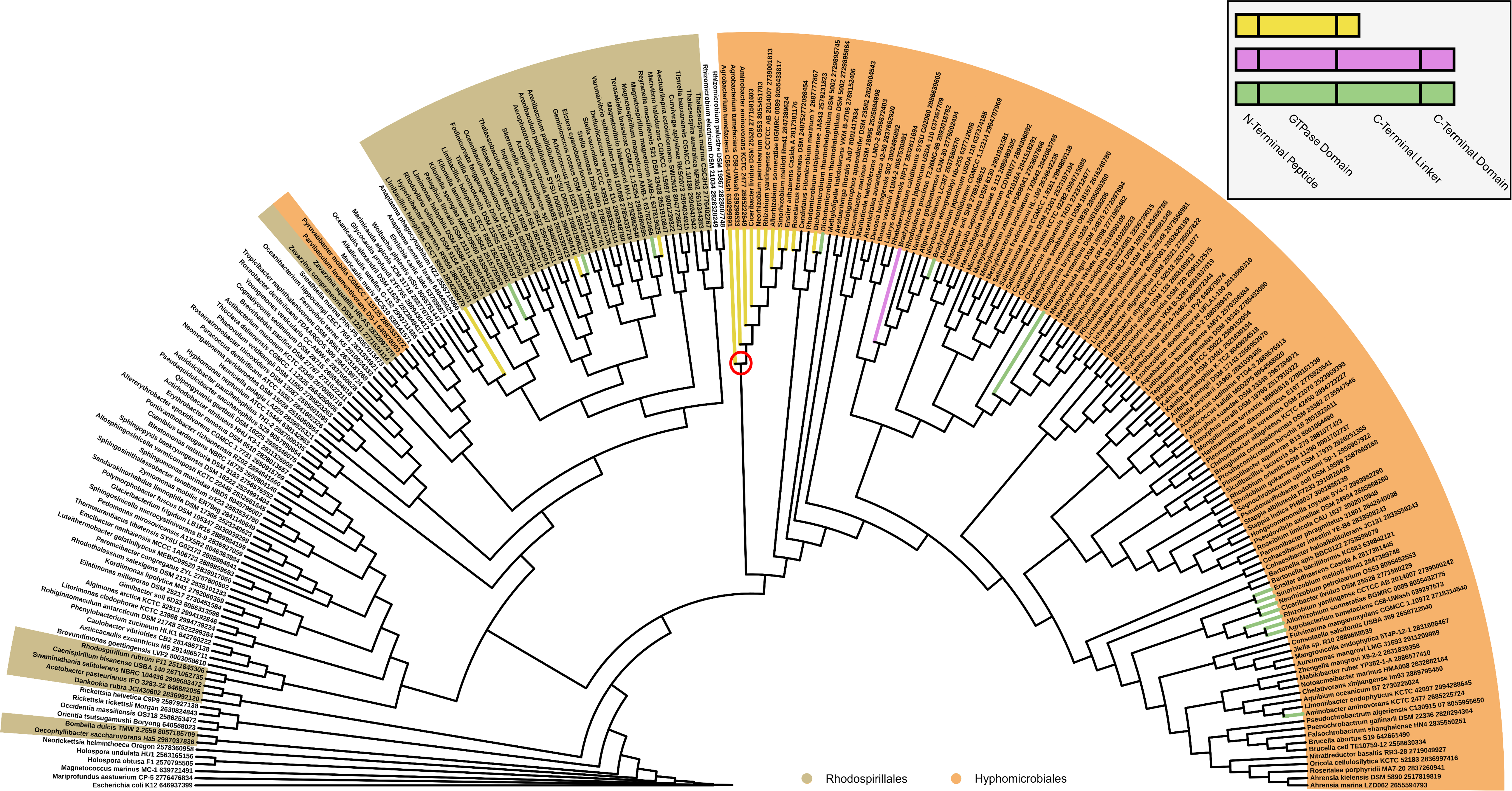
FtsZ duplications have arisen multiple times in Alphaproteobacterial class. GTPase domain tree of select Alphaproteobacteria reveals gene duplications across several orders of Alphaproteobacteria. Branches in black indicate that species that have a single copy of FtsZ. Colored branches indicate species with FtsZ duplications. Green branches lead to canonical FtsZ while yellow and purple branches indicate duplicated FtsZ paralogs. Representative domain structures for canonical (green) and duplicate (yellow & purple) FtsZ copies are shown in the grey box at the top-right. *Bradyrhizobium japonicum* is the only duplicate in the Hyphomicrobiales order that possesses all four structural motifs for FtsZ. Red circle indicates a node comprised solely of duplicated FtsZ paralogs. Color coded names indicate the Alphaproteobacterial order of the organisms containing the FtsZ protein.

Remarkably, duplicate *ftsZ* sequences can be found outside of the *dcw* operon, but most of them are truncated and missing one or more of the C-terminal motifs (Vaughan et al. 2004). Within the Alphaproteobacterial class, Vaughan et al. described six duplicates containing NTP, GTPase Domain, and partial CTL motifs. For the five duplicates from the Hyphomicrobiales order, they reported differences between the truncated duplicate *ftsZ* genes from the *Rhizobiaceae/Phyllobacteraceae* families and the *Bradyrhizobium japonicum* genome in the *Nitrobacteraceae* family, which contains two full length *ftsZ* sequences (Vaughan et al. 2004). Since this description, genome sequencing efforts have extended the available dataset across the Alphaproteobacterial class. In this work, we expanded the analysis of *ftsZ* duplications to each described genus in the Alphaproteobacterial class as well as 831 species within the Hyphomicrobiales order. This analysis provided further evidence that two independent *ftsZ* duplications occurred within the Hyphomicrobiales order of the Alphaproteobacterial class, resulting in distinct *Rhizobiaceae*/*Phyllobacteraceae* and *Nitrobacteraceae* lineages. Mapping the unique substitution patterns in these two groups to structural models suggests positive selection for functions distinct from canonical FtsZ. We explored possible functions of the *ftsZ* duplications in the *Rhizobiaceae*/*Phyllobacteraceae* group by manipulating *ftsZ*_1_ and *ftsZ*_3_ genes in *Agrobacterium tumefaciens*. Remarkably, overexpression of FtsZ_1_ resulted in a complete block in cell division, producing large, round cells indicative of misregulation of peptidoglycan biosynthesis. Consistent with this possibility, we found that a hyperactive allele of *ftsW,* the monofunctional transglycosylase for septal peptidoglycan synthesis, partially protected against FtsZ_1_ overexpression. Overall, these results indicate that the truncated FtsZ_1_ paralog in *A. tumefaciens* contributes to regulation of division processes such as septal peptidoglycan synthesis. Furthermore, since *ftsZ* duplications are maintained disproportionately in Alphaproteobacteria associated with plants, we speculate that FtsZ paralogs may function to fine tune cell division *in planta* or related environments.

## Materials and Methods

### Bacterial strains, plasmids, and growth conditions

All bacterial strains and plasmids used in this study are provided in the Strain List found in File S2. *Agrobacterium tumefaciens* was grown in *A. tumefaciens* glucose & nitrogen (ATGN) minimal media at 28°C shaking (Tempé et al. 1977; Morton & Fuqua, 2012). For plasmid selection, kanamycin was used at a working concentration of 300 µg/mL for *A. tumefaciens.* When culturing *Escherichia coli*, lysogeny broth (LB) medium was used, and cells were grown at 37°C shaking (Bertani, 1951). When appropriate, 50 µg/mL of kanamycin was added.

For spotting assays, indicated media was prepared with 1.5% agar. The agar plates included ATGN minimal media in the main-text figures as well as the following media for Figure S3 in File S1: ATSN minimal media, LB, LB0 (which follows the same recipe of LB without NaCl). When appropriate, ATGN was supplemented with ampicillin, cephalexin, and aztreonam to a final concentration of 10 µg/mL, cefsulodin to 25 µg/mL, acetosyringone to 200 µM, kanamycin to 300 µg/mL, or 1 mM isopropyl β-D-1-thio galactopyranoside (IPTG).

### Construction of plasmids and strains

FtsZ_1_ and FtsZ_3_ were PCR amplified with 20 base pairs of overlap, with regards to the amplified pSRKKM Ptac backbone, using primers listed in the Plasmid & Primer List in the File S2. The PCR products were joined using Gibson Assembly to construct the plasmids listed in the Plasmid & Primer List in the File S2. Each plasmid was transformed into *E. coli* DH5α strains following the New England Biolabs standard protocol for heat-shock transformations. Plasmids were maintained in *E. coli* DH5α and were miniprepped and sequence-verified (Plasmidsaurus). Successful constructs were electroporated into *E. coli* S17-1 λpir, which was subsequently used to transfer the plasmid into *A. tumefaciens* wildtype or FtsW F137L (FtsW*) backgrounds. The mating procedure occurs by mixing cell pellets of overnight S17-1 λpir+ vector and *A. tumefaciens* cultures, each resuspended in 50 µL and placed onto sterile filter paper on LB IPTG plates. After 24 hours of incubation at 28°C, cells were collected and serially diluted onto ATGN Kan300 µg/mL plates: which *E. coli* cannot grow on.

### Alphaproteobacterial FtsZ duplication phylogenetic tree construction

Using the Joint Genome Institute (JGI) database, at least one representative from each genus within the Alphaproteobacteria was identified (Grigoriev et al. 2012; Nordberg et al. 2014). Within this set of genera, we prioritized those with genomes that were finished or permanent drafts. The imbedded NCBI protein-protein BLAST function was used to identify FtsZ copies within each genome (Altschul et al. 1990). First, we conducted a BLAST search with FtsZ_AT_ from *Agrobacterium tumefaciens.* We repeated BLAST searches using FtsZ sequences from *Sphingosinicella microcystinivorans, Anaplamsa phagocytophilum, Ehrlichia canis, Magnetococcus marinus,* and FtsZ_1_ from *Agrobacterium tumefaciens* as a query. No additional FtsZ sequences were identified with the additional BLAST searches suggesting that our searches identified *ftsZ* paralogs within these genera.

Once the FtsZ sequences were identified, MEGA11 was utilized to align all FtsZ sequences (Tamura et al. 2021). For this process, the imbedded CLUSTAL alignment tool was used. Then, FtsZ_AT_ was used as a guide to trim all sequences down to the GTPase domain. Finally, the best model for substitutions was estimated to be LG+G+I using MEGA11. Next, we uploaded the aligned, truncated FtsZ file to BEAST v1.10.4 and used the substitution model LG+G+I. For the “Tree Prior” setting, we selected Speciation: Yule Process (Yang, 1994; Le & Gascuel, 2008; Gernhard, 2008; Drummond et al. 2012; Suchard et al. 2018). When appropriate, the seed 1708101569949 was used. For the MCMC settings, the length of chain was set to 10,000,000 with screening every 1,000. After running through the 10,000,000 iterations on BEAST, the Maximum Clade Credibility tree was constructed with TreeAnnotator: part of the BEAST package. We followed the default suggestion of using 10% of all states calculated: 1,000,000. The Maximum Clade Credibility tree was visualized with ITol (Letunic & Bork, 2021). Canonical FtsZ sequences were identified using the built-in genome viewer of JGI/IMG. Copies found in the *division and cell wall* (*dcw*) operon were identified as the canonical FtsZ copy.

### Cell viability spotting assay

Cells were inoculated into 1 mL of ATGN and incubated for overnight growth of 16-21 hours. Each strain was diluted into a new culture tube to result in an OD_600_ = 0.1. Afterwards, 4-6 hours of outgrowth allowed for cultures to reach exponential phase growth (OD_600_ = 0.4-0.6). Each strain was then diluted to OD_600_ = 0.1 before being serially diluted in 10-fold increments from series 10^0^ through 10^-7^. After diluting, 3 μL of each dilution was pipetted onto 1.5% ATGN agar plates. Plates were allowed to dry for 1 hour at room-temperature and then were transferred to 28°C for 72 hours. Plates were imaged with the BioRad ChemiDoc MP imager.

### Microscopy and quantitative image analysis

To prepare for microscopy, strains were grown to exponential phase as described for the viability assay. For phase microscopy, 0.5 μL of cells were spotted onto 1.75% ATGN agarose pads and imaged using an inverted Nikon Eclipse TiE with a QImaging Rolera em-c^2^ 1K EMCCD camera and Nikon Elements Imaging Software. For overexpression experiments, 1 mM IPTG was added to the liquid medium and bacterial populations were imaged by spotting the cultures onto pads at specific timepoints. For timelapse microscopy, 1 mM IPTG was instead added to agarose pads and images were acquired every 10 minutes for a set field of views. Phase microscopy images were converted to binary masks using ImageJ (Rueden et al. 2017). Binary masks were processed using CellTool to measure the area of each cell outline following the CellTool tutorial (Pincus & Theriot, 2007). Area measurements were visualized using R-Studio (R Core Team, 2023).

### Hidden Markov and PyMOL modeling

FtsZ and FtsZ GTPase duplications were identified and collected from genomes on JGI, as described above, but every genome was selected within the Hyphomicrobiales order shared between JGI and the List of Prokaryotic names with Standing in Nomenclature (Table S2, File S2; Parte et al. 2020). Protein sequences from *Nitrobacteraceae* (canonical n = 36, duplicate n = 38), *Phyllobacteraceae* (canonical n = 80, duplicate n = 84), and *Rhizobiaceae* (canonical n = 142, duplicate n = 157) families were aligned to FtsZ GTPase domains from *E. coli* (IMG ID 2978828590) and *A. tumefaciens* (NP_532761.1) as separate groups using Jalview (Troshin et al. 2011). The GTPase domain was defined as the first 315 amino acids of the *E. coli* FtsZ GTPase domain. Only sequences with unambiguous alignment and full coverage of the GTPase domain were included in this analysis. When necessary, the N-termini of sequences were trimmed to the positions of the conserved 15 amino acids of canonical FtsZ from Hyphomicrobiales NTP (Figure 3b). WebLogo3 was used to plot the amino acid distribution at each position of the GTPase domain (Suchard et al. 2018). Structural models were generated with PyMol using 6UNX (PDB) as a scaffold and color-coding the conservation patterns using the HMMs that maintained *E. coli* amino acid numbering (The PyMOL Molecular Graphics System, Version 1.2r3pre, Schrödinger, LLC.).

**Figure 2.**
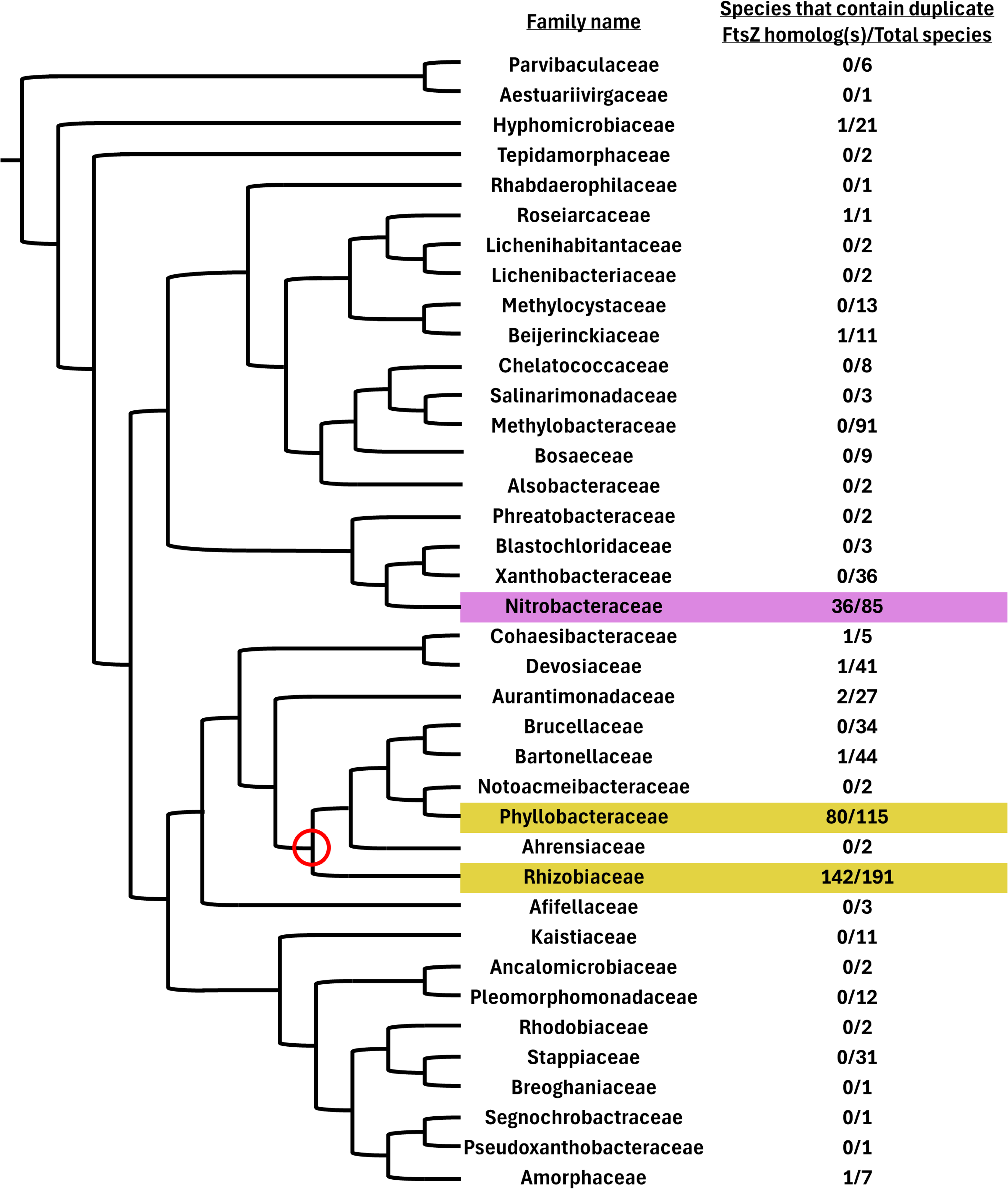
FtsZ duplications are present, primarily, in three families of the Hyphomicrobiales order. (a) Using the Joint Genome Institute/Integrated Microbial Genomes & Microbiomes database, each species detailed within the Hyphomicrobiales order, as annotated by the List of Prokaryotic names with Standing in Nomenclature, was assessed for the presence or absence of FtsZ duplications with protein similarity searches and synteny comparisons. FtsZ sequences identified were grouped by family without regard of multiple duplications. Cladogram was drawn with no evolutionary data but was based off previous work (diCenzo et al. 2024). Red circle indicates the last shared node between Rhizobiaceae and Phyllobacteraceae families.

**Figure 3.**
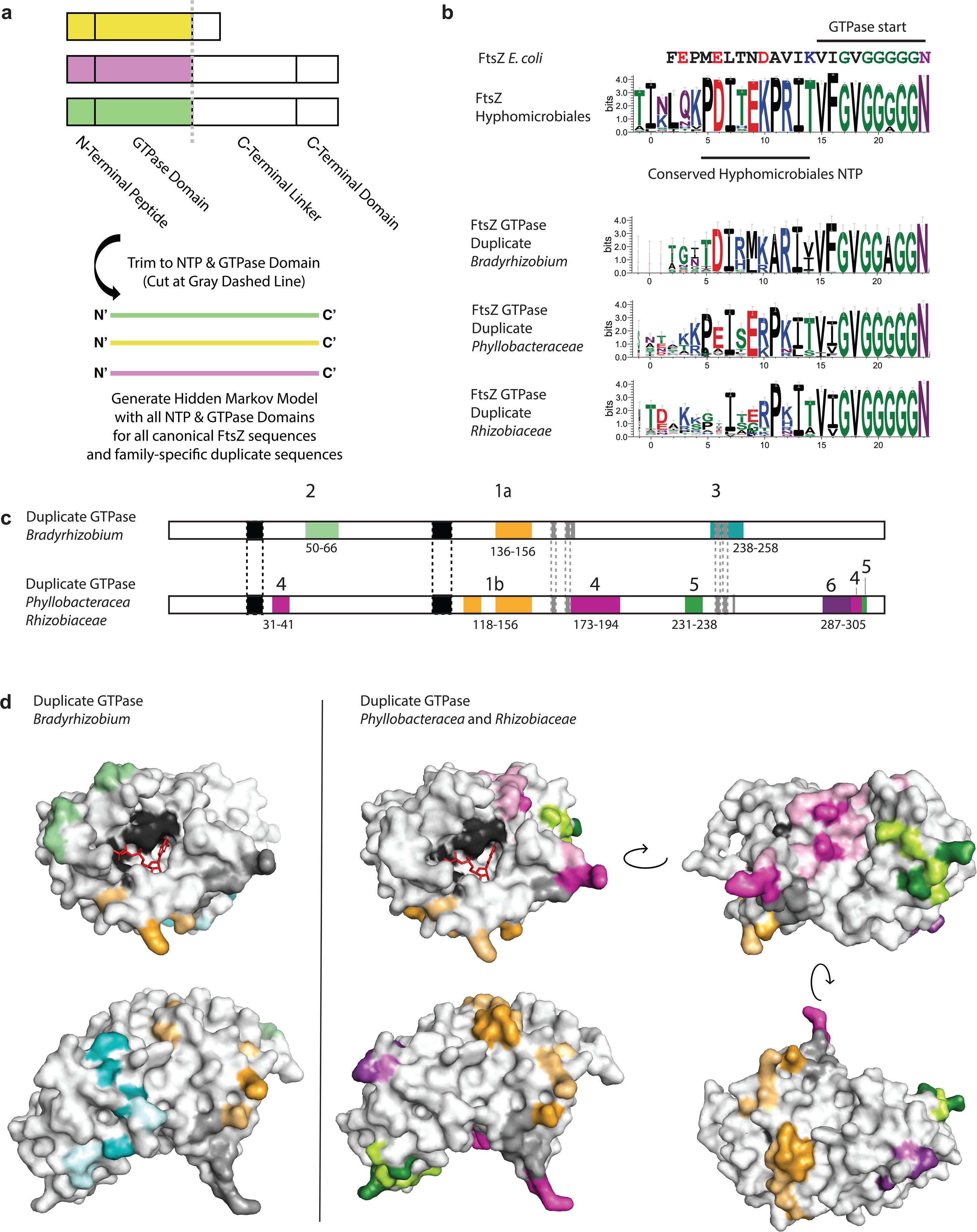
FtsZ GTPase duplicates are conserved within three Hyphomicrobiales families and constitute two distinct lineages. (a) Schematics of FtsZ and its paralogous GTPase domain duplications and explanation of approach. (b) Hidden Markov Model (HMM) logos indicating the consensus protein sequences of the leader regions of Hyphomicrobiales FtsZ (n = 260 sequences) and the duplications identified in the *Bradyrhizobium* (n = 38 sequences), *Phyllobacteraceae* (n = 82 sequences), and *Rhizobiaceae* families (n = 152 sequences) using WebLogo 3 (Suchard et al. 2018). Amino acids are color coded according to chemical properties, with uncharged polar residues in green, neutral residues in purple, basic residues in blue, acidic residues in red, and hydrophobic residues in black. The height of each letter is proportional to the relative frequency of a given identity, and the height of the stack indicates the sequence conservation at that position. E. coli FtsZ numbering is used for ease of comparison. (c) Schematic summary of HMM analysis of the entire duplicated GTPase domains. Because *Phyllobacteracea* and *Rhizobiaceae* share the same HMM patterns, they are shown as one schematic. Black regions are large, conserved regions that form the GTP-binding catalytic site (Löwe & Amos, 1998). Gray regions are putative FzlA binding residues estimated from *C. crescentus* (Payne et al. 2024). Dashed lines between the two sequences indicate conservation across all three families. Numbered, colored boxes indicate remodeled regions of interest that have been mapped to the structural models in (d) and (e). Full length HMMs are provided in Figure S2, File S1. Numbering is based on *E. coli* FtsZ. (d) Remodeled regions from the HHM of the *Bradyrhizobium* GTPase duplicate mapped onto *E. coli* FtsZ (PDB 6UNX). Colored regions correspond to the remodeled regions indicated in (c) with dark shading indicating a conserved substitution with a chemical change from canonical FtsZ GTPase and light shading indicating loss of conservation from canonical FtsZ GTPase (see Figure S2, File S1 for HHM and residue identities). GTP is shown in red. (e) Remodeled regions from the HHM of the *Phyllobacteracea* and *Rhizobiaceae* GTPase duplicate, as in (d).

## Results and Discussion

### Duplicate FtsZ proteins are distributed throughout the Hyphomicrobiales Order

Several FtsZ duplicates have previously been identified within the Alphaproteobacterial class including those found in *Agrobacterium tumefaciens, Sinorhizobium meliloti, Bradyrhizobium japonicum,* and *Magnetospirillum magneticum* (Vaughan et al. 2004). To better understand the distribution and function of these FtsZ duplicates, we took advantage of the rapidly growing number of publicly available whole-genome-sequences to generate a FtsZ GTPase domain tree using representatives from each genus across the Alphaproteobacterial class. FtsZ protein sequences were identified through iterative NCBI BLAST using the FtsZ sequences from *A. tumefaciens, Sphingosinicella microcystinivorans, Ehrlichia canis, Magnetococcus marinus,* and the FtsZ_1_ sequence from *A. tumefaciens* (Table S1, File S2). Each BLAST iteration resulted in the same results, indicating that the identified FtsZ duplications share enough conservation with the canonical FtsZ sequences to allow differentiation from other GTPase domain-containing genes. For species that possessed multiple hits, the FtsZ copy encoded in the *dcw* operon was identified as the canonical sequence while those outside of this operon were classified as duplicates.

Known duplications across *A. tumefaciens*, *S. meliloti,* and *B. japonicum* were observed (Vaughan et al. 2004), as well as previously unidentified FtsZ duplicates in *Limibacillus halophilus, Arenibaculum pallidiluteum, Aminobacter aminovorans, Ciceribacter lividus, Neorhizobium petrolearium, Rhizobium yantingense, Allorhizobium sonneratiae, Ensifer adhaerens, Roseiarcus fermentans,* and *Dichotomicrobium thermohalophilum* (Figure 1; Figure S1, File S1). Notably, the duplicate identified in *L. halophilus,* as well as the previously described duplicate in *B. japonicum*, were the only duplicates that possessed a CTD while all other duplicates ended with GTPase domains or minimal CTLs (Figure 1, grey box). Aside from this, two patterns of FtsZ duplications were readily observed: first, some FtsZ duplicates were most closely related to the canonical FtsZ GTPase domain from the same organism. This was observed in the Rhodospirillales order (*A. pallidiluterum* and *M. magneticum*) as well as the Hyphomicrobiales order (*D. thermohalophilum*) (Figure 1; Figure S1, File S1). This pattern is consistent with recent gene duplications giving rise to the secondary copies of FtsZ within each species. Second, the majority of the identified FtsZ duplicates were instead more closely related to duplicate FtsZ GTPase domains from other species than to the canonical FtsZ in the same species. A clade comprised solely of FtsZ duplicates in the Hyphomicrobiales order consisted of *A. tumefaciens, A. aminovorans, C. lividus, N. petrolearium, R. yantigense, A. sonneratiae, S. meliloti,* and *E. adhaerens* (marked by a red circle at the base of the clade in Figure 1 & Figure S1, File S1). The formation of this clade within the *Rhizobiaceae* and *Phyllobacteraceae* families suggests that these FtsZ GTPase duplications are potentially orthologs of each other with a distinct evolutionary trajectory from canonical FtsZ GTPase domains. Vaughan *et al*. also identified duplications in each of these families and our observations are consistent (Vaughan et al. 2004).

To better understand FtsZ duplication patterns within the Hyphomicrobiales, we completed a fine-scale analysis of the distribution of FtsZ duplicates within this order. Consistent with observations in Figure 1, we found that FtsZ duplications within *Roseiarcaceae* and *Hyphomicrobiaceae* are rare (Figure 2; Table S2, File S2). Interestingly, the duplicate FtsZ GTPase domain from the *Roseiarcaceae* member, *R. fermentans*, falls just outside of the duplicate clade in Figure 1 while also being separated from its respective canonical sequence. As for the representative from *Hyphomicrobiaceae, D. thermohalophilum,* its duplicate and canonical sequences pair together (Figure 1; Figure S1, File S1). However, these observations could be due to the relatively limited number of genome sequences published for these families and further analysis, as new data becomes available, will be necessary.

Retention of a truncated FtsZ duplicate occurs at a very high frequency (∼70%) in the *Rhizobiaceae* and *Phyllobacteraceae* families (Figure 2; Table S2, File S2). Given that these duplicates are also more closely related to each other than the canonical FtsZ in the same species (Figure 1), this pattern is consistent with an ancient acquisition of a duplicate FtsZ GTPase prior to the divergence of these families. We identified another radiation of a FtsZ duplication within the *Nitrobacteraceae,* with over 40% of the sequenced species retaining a duplicate, but this was constrained to the genus *Bradyrhizobium* (Figure 2; Table S2, File S2). Therefore, the size of this cluster of duplications is due to the extensive sampling of this genus for genomic sequencing. Moreover, all *Bradyrhizobium* duplicates possessed the four canonical FtsZ domains. This contrasts with the *Rhizobiaceae* and *Phyllobacteraceae* duplications where nearly all the duplicates are truncated to the NTP, GTPase Domain, and short CTLs (Figure 1, upper right inset). Hereafter, we refer to the *Nitrobacteraceae* duplicates as the *Bradyrhizobium* duplicates since all duplications are found within that genus. Based on these observations, we next used a bioinformatic approach to explore the similarities and differences between the *Rhizobiaceae*, *Phyllobacteraceae*, and *Bradyrhizobium* canonical and duplicate FtsZ GTPases.

### Two independent FtsZ GTPase duplications have arisen and have been maintained in the Hyphomicrobiales Order

To understand the relationship of the Hyphomicrobiales duplicate FtsZ proteins with their canonical FtsZ counterparts and with each other, we created amino acid consensus sequences for the canonical and duplicate FtsZ GTPase domains (Figure S2, File S1). Because we expected to see strict conservation of residues required for the function of the domain, comparison of these consensus sequences could pinpoint which regions of the GTPase domain are under purifying selection in the duplicates (Figure 3a). Residues with high conservation in the duplicate FtsZ GTPases that are chemically distinct from conserved residues in the canonical FtsZ GTPase domain could be under selection for new function. In contrast, positions with high variability in the duplicate FtsZ GTPase that are strictly conserved in the canonical FtsZ GTPase domain are no longer under selection for the original structure and/or function of the FtsZ GTPase domain. Identifying regions with the combination of these patterns would suggest which portions of the duplicate FtsZ GTPase domain have been remodeled for a function distinct from that of the canonical FtsZ protein.

As expected, the canonical FtsZ GTPase domains exhibit high conservation with the *E. coli* FtsZ GTPase domain except for the NTP (Figure 3b). The NTP has extensive variation across the bacterial domain, varying from as few as six residues in some firmicutes to as many as seventy-seven residues in actinomycetes (Vaughan et al. 2004). This variation could underlie family-specific differences with interacting regulatory partners. For instance, the first 32 residues of *E. coli* FtsZ have been reported to interact with FtsZ-inhibiting proteins YeeV (Tan et al. 2010) and SulA (Huang et al. 1996). Furthermore, the conserved 51 residue NTP of heterocyst-forming cyanobacteria has been shown to contribute to the polymerization behavior and binding interactions with the essential divisome component SepF in *Anabaena sp. PCC 7120* (Corrales-Guerrero 2018). This region ranges from 7-44 amino acids in the Alphaproteobacteria (Vaughan et al. 2004) although no regulatory function has been reported for any members. The portion of the NTP proximal to the start of the GTPase domain, PDITEKPR, is invariant among all the canonical Hyphomicrobiales sequences with the preceding six N-terminal residues exhibiting general conservation with some substitutions (Figure 3b). The strict maintenance of this region within this order suggests that it may behave as a scaffold for protein-protein interactions.

In comparison to the strict conservation in the canonical FtsZ sequences, the duplicate NTPs exhibit diminishing conservation by sequence identity or similarity (Figure 3b). In particular, the first six or seven residues are highly variable, constituting the largest remodeled region in all the HMMs (Figure S2, File S1), followed by mostly conservative substitutions in the PDITEKPR sequence. The duplicate FtsZ GTPases from the *Rhizobiaceae* family exhibit the most variation and loss of conservation in the NTP. Overall, the loss of strict conservation of the NTP in the duplicate FtsZ GTPase domains indicates that this region is under different evolutionary constraints than the canonical FtsZ GTPase domain. Thus, we hypothesize that the canonical and duplicate FtsZ proteins are not regulated at the N-terminus by the same regulatory networks in the

### Rhizobiaceae and Phyllobacteraceae families

In stark contrast to the variation seen in the NTP region, the remainder of the duplicate FtsZ GTPase domains strictly conserve the GTP-binding pocket (Figure 3c black boxes and 3de black residues; Figure S2, File S1; Löwe & Amos, 1998) and residues predicted to bind the Alphaproteobacterial divisome protein FzlA (Figure 3c gray boxes and 3de gray residues; Figure S2, File S1; Payne et al. 2024). This suggests that the duplicate GTPases are still active and could overlap in function with the canonical FtsZ GTPase domain in the Z-ring. There is support for this in *A. tumefaciens,* where the duplicate FtsZ_1_ likely forms functional co-polymers with FtsZ_AT_ (Howell et al. 2019). The co-polymers form similar structures and maintain GTP hydrolysis rates comparable to homopolymers of FtsZ_AT_ *in vitro*. *In vivo*, localization of FtsZ_1_ to mid-cell is dependent on the presence of FtsZ_AT_ (Howell et al. 2019). These observations suggest that the FtsZ duplicates may regulate cell division when incorporated into the Z-ring. It is intriguing to speculate that the conditional formation of FtsZ co-polymers may tune cell division by promoting or inhibiting Z-ring dynamics in response to differing environmental conditions.

When compared to the canonical FtsZ consensus sequence, the consensus sequences of the duplicate FtsZ GTPase domains from *Phyllobacteraceae* and *Rhizobiaceae* share the same remodeling pattern (Figure 3c; Figure S2, File S1), whereas the duplicate FtsZ GTPase domain from the *Bradyrhizobium* group exhibits a distinctly different pattern with fewer substitutions. We mapped these positions to structural models of *E. coli* FtsZ GTPase domain to understand their spatial relationship to each other and to conserved regions known to interact with GTP or regulatory protein FzlA (Figure 3de). Figure 3c summarizes six regions of denser remodeling of proximal surface residues to illustrate the two different substitution patterns (see Figure S2, File S1 for entire consensus sequences with identified regions). These remodeled regions were defined both by positions with invariant substitutions that change the chemical nature of the residue (darker hue) and positions with increased variation or relaxed conservation (lighter hue). The remodeling of these regions could modify the behavior of this duplicate FtsZ GTPase domain from the canonical FtsZ GTPase domain by facilitating interactions with different binding partners.

Overall, the remodeling patterns suggest that the *Phylobacteraceae* and *Rhizobiaceae* duplicate FtsZ proteins share a common ancestor while the *Bradyrhizobium* duplication is an independent, more recent event. The lower number of substitutions in the *Bradyrhizobium* duplicate consensus sequence is expected for a duplication occurring more recently within a single genus. The clear split between the two lineages of duplicate FtsZ proteins agrees with the known phylogenetic relationships of these families (Figure 2) and previous observations from studies with fewer genomes (Vaughan et al. 2004). However, this observation also suggests that the duplication, while maintained in the *Phyllobacteraceae* and *Rhizobiaceae* families, has been lost in four other Hyphomicrobiales families that share the same last common ancestor (Figure 2, red circle). The Hyphomicrobiales exhibit extensive lifestyle diversity (Williams et al. 2022). Because the *Phylobacteraceae* and *Rhizobiaceae* families are known for soil-based lifestyles and plant interactions, while *Brucella* and *Bartonella* are families of animal pathogens, we wondered if the lifestyles of the families lacking the duplicate FtsZ GTPase were distinctly different. Within the families *Ahrensiaceae* and *Notoacmeibacteracea*, we found that most species were originally isolated from marine environments (Table S3, File S2; Jung et al. 2012; Liu et al. 2016; Huang et al. 2017; Yan & Tuo, 2019). Likewise, for genera that have lost the duplication in the *Phyllobacteraceae* and *Rhizobiaceae* families, most were also isolated from marine environments or hosts (Table S3, File S2; Roh et al. 2008; Kämpfer et al. 2015; Li et al. 2016; Hyeon et al. 2017; Tóth et al. 2017; Cao et al. 2020; Song et al. 2023; Wang et al. 2023). Together these observations support the possibility that an ancient duplication of FtsZ has been maintained in the *Phyllobacteraceae* and *Rhizobiaceae* lineages as an adaptation for specific terrestrial plant-associated or soil lifestyles. It is possible that the duplication in *Bradyrhizobium* arose convergently under the similar selective pressures, however its distinct substitution pattern and limited distribution within a single genus makes it impossible to know whether it is being retained for similar function.

### FtsZ_1_ overexpression inhibits cell division

Overall, the maintenance of the duplicate FtsZ within the *Rhizobiaceae* and *Phyllobacteraceae* lineages suggests that the duplicate FtsZ proteins may be under selection for a distinct function, such as an alternative GTPase in the Z-ring to adapt to stresses experienced in the soil environment. We used *A. tumefaciens* as a model to explore possible functions of FtsZ duplicates in a soil dwelling microbe with the ability to invade a plant host. *Agrobacterium tumefaciens* possesses two FtsZ duplicates, FtsZ_1_ and FtsZ_3_, that are non-essential in standard laboratory conditions (Howell et al. 2019). Here, we exposed deletion mutants to a larger array of conditions, including host invasion and virulence. FtsZ_1_ and FtsZ_3_ were dispensable for viability in all tested conditions (Figure S3, File S1). Next, we hypothesized that FtsZ duplicates may function as negative regulators of cell division. To test this hypothesis, we overexpressed FtsZ_1_ and FtsZ_3_ and found that FtsZ_1_ overexpression resulted in a significant, 5-log decrease in viability (Figure 4a). This suggests that FtsZ_1_ inhibits an essential activity when expressed at high levels. In contrast, FtsZ_3_ overexpression did not impact viability (Figure 4a), morphology, or frequency of cell division (Figure S4, File S1).

**Figure 4.**
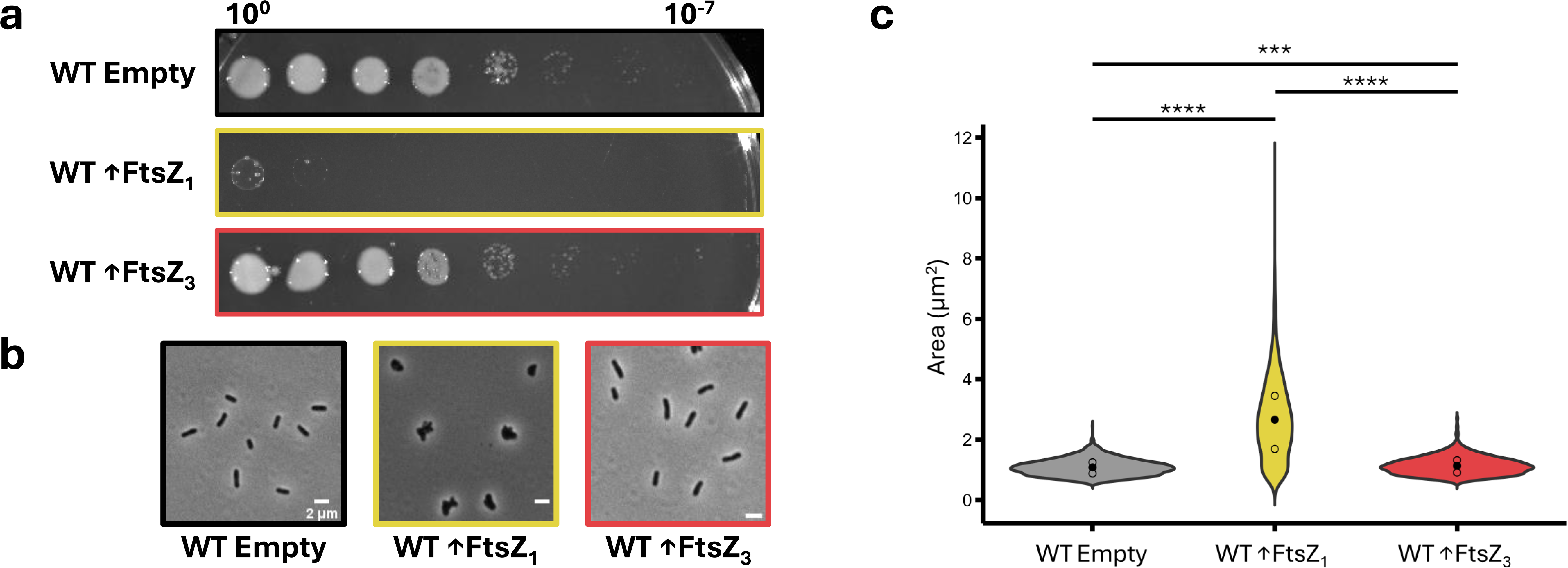
Overexpressing FtsZ_1_ results in a dramatic morphology and viability defect. (a) Spotting assays of *A. tumefaciens* in the wildtype background with pSRKKM Ptac: empty vector (Empty), FtsZ_1_and FtsZ_3_. When grown on ATGN Kan300 µg/mL 1mM IPTG, the protein listed to the left is expressed from the plasmid. (b) Phase contrast microscopy of strains from previous panel after 16 hours of induction. (c) Area measurements (µm^2^) after expressing the Empty Vector, FtsZ_1_, or FtsZ_3_ for 16 hours. Black dots indicate the mean for each sample while empty dots represent the 1st and 3rd quartile (n=1844, 3394, 3104). Comparing each mean, via Kruskal-Wallis non- parametric ANOVA test and a Dunn’s post hoc test, p-values are represented by *, p < 0.05; **, p < 0.01; ***, p < 0.001; ****, p < 0.0001.

Because duplicate FtsZ GTPases might regulate the essential activity of FtsZ_AT_ during division, we used quantitative image analysis to assess morphological changes 16 hours after induction of FtsZ_1_ or FtsZ_3_ (Figure 4bc). In agreement with the viability effect, FtsZ_1_ overexpression resulted in dramatic changes in morphology. Cells overexpressing FtsZ_1_ lost their rod-shaped morphology and appeared to swell from mid-cell while also forming ectopic poles (Figure 4b). Unexpectedly, this phenotype is not consistent with inhibiting FtsZ _AT_ activity; when FtsZ_AT_ is depleted, cells fail to terminate polar growth, and growth-active poles undergo tip splitting events leading to the formation of elongated cells which accumulate growth poles and, ultimately, lyse (Howell et al. 2019). The depletion of FtsZ_AT_ prevents the establishment of FtsZ rings and termination polar growth. In contrast, the phenotype associated with FtsZ_1_ overexpression suggests that FtsZ rings form and polar growth terminates properly; however, the mid-cell bulges and ectopic pole formation suggest that septal peptidoglycan synthesis is misregulated and polar growth may commence despite the block in cell division.

### Hyperactive FtsW partially bypasses the effects of FtsZ_1_ overexpression

We hypothesized that if FtsZ_1_ overexpression impaired septal peptidoglycan biosynthesis pathways, hyperactive alleles of FtsW and/or FtsI would protect *A. tumefaciens* from FtsZ_1_ overexpression. These hyperactive backgrounds have been shown to rescue other strains with impaired septal peptidoglycan synthesis by bypassing a late checkpoint in division (Modell et al. 2014; Lariviere et al. 2019; Li et al. 2021; Attaibi & Blaauwen, 2022; Payne et al. 2024). Therefore, we tested if FtsW*, a hyperactive allele of FtsW (F137L) could rescue the toxic phenotype caused by FtsZ_1_ overexpression. Indeed, the FtsW* background restored viability of cells overexpressing FtsZ_1_ to that of the parent strain (Figure 5a). Quantitative image analysis indicated that the presence of hyperactive FtsW* reduced the cell area in both the wildtype and FtsZ_1_ overexpression strains (Figure 5bc), presumably due to a lessened activation requirement for the septal peptidoglycan synthase. Despite this rescue, morphological defects persisted in these cells. Image analysis of cells 16 hours post-induction revealed that most cells overexpressing FtsZ_1_ still exhibited swelling and ectopic pole formation despite restored viability (Figure 5b). In conclusion, the hyperactivation of septal peptidoglycan synthesis was sufficient to promote cell division and rescue viability, but without resolving the morphological defect.

**Figure 5.**
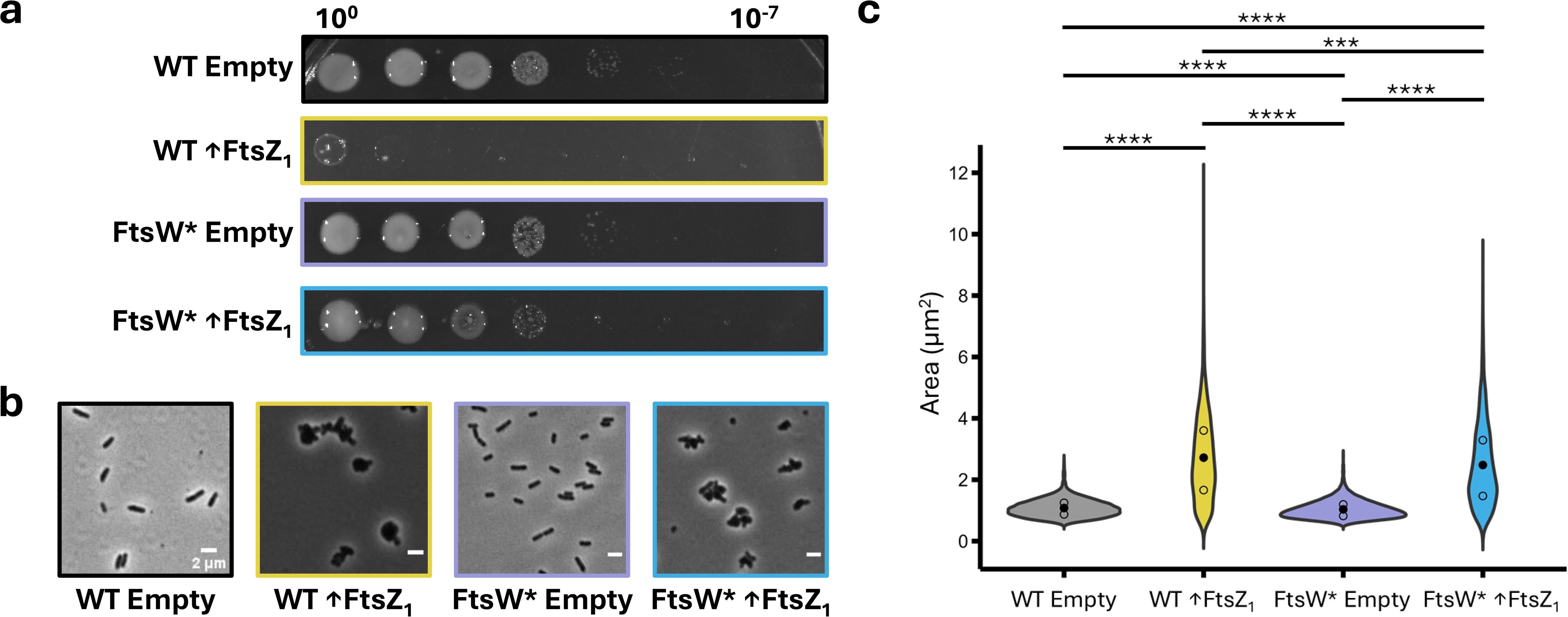
A hyperactive FtsW allele (FtsW*) partially resolves the phenotype of FtsZ_1_ overexpression. (a) Spotting assays of *A. tumefaciens* in the wildtype background with pSRKKM Ptac: empty vector and FtsZ_1_ as well as the FtsW* background with pSRKKM Ptac: Empty and FtsZ_1_. When grown on ATGN Kan300 µg/mL 1mM IPTG, the protein listed to the left is expressed from the plasmid. (b) Phase contrast microscopy of strains from previous panel after 16 hours of induction. (c) Area measurements (µm^2^) after 16 hours of induction. Black dots indicate the mean for each sample while empty dots represent the 1^st^ and 3^rd^ quartile (n= 3302, 3108, 3110, 1947). Comparing each mean, via Kruskal-Wallis non-parametric ANOVA test and a Dunn’s post hoc test, *p-values* are represented by *, p < 0.05; **, p < 0.01; ***, p < 0.001; ****, p < 0.0001.

The observation that pleomorphic cells have wildtype-like viability is counterintuitive. We used timelapse microscopy to better understand this complex phenotype and understand how these cells maintain viability during FtsZ_1_ overexpression in the FtsW* background we used timelapse microscopy. For this experiment, instead of growing the cells for 16 hours after inducing overexpression of FtsZ_1_, cells were induced on agarose pads and imaged over time to capture morphological and division defects in real time. Consistent WT rod-shaped morphology and division events occurred in cells containing the empty vector (Figure 6a). Wildtype cells overexpressing FtsZ_1_ were inhibited in cell division within 2 hours and mid-cell swelling and/or ectopic pole formation was evident by 4 hours, leading to loss of proper rod-shaped morphology (Figure 6a). In comparison, the FtsW* background restored cell division events throughout FtsZ_1_ overexpression for most cells up to 4 to 6 hours. We used the timelapse data to quantify the frequency of cell septation, division, and lysis in each strain (Figure 6b). When FtsZ_1_ was overexpressed in a wildtype background, only ∼20% of cells divided; however, in the FtsW* background ∼70% of cells divided (Figure 6b). These results confirm our previous observations of whole populations 16 hours post-induction and further support that overexpression of FtsZ_1_ results in a block in cell division that can be overcome with the hyperactive FtsW* background.

**Figure 6.**
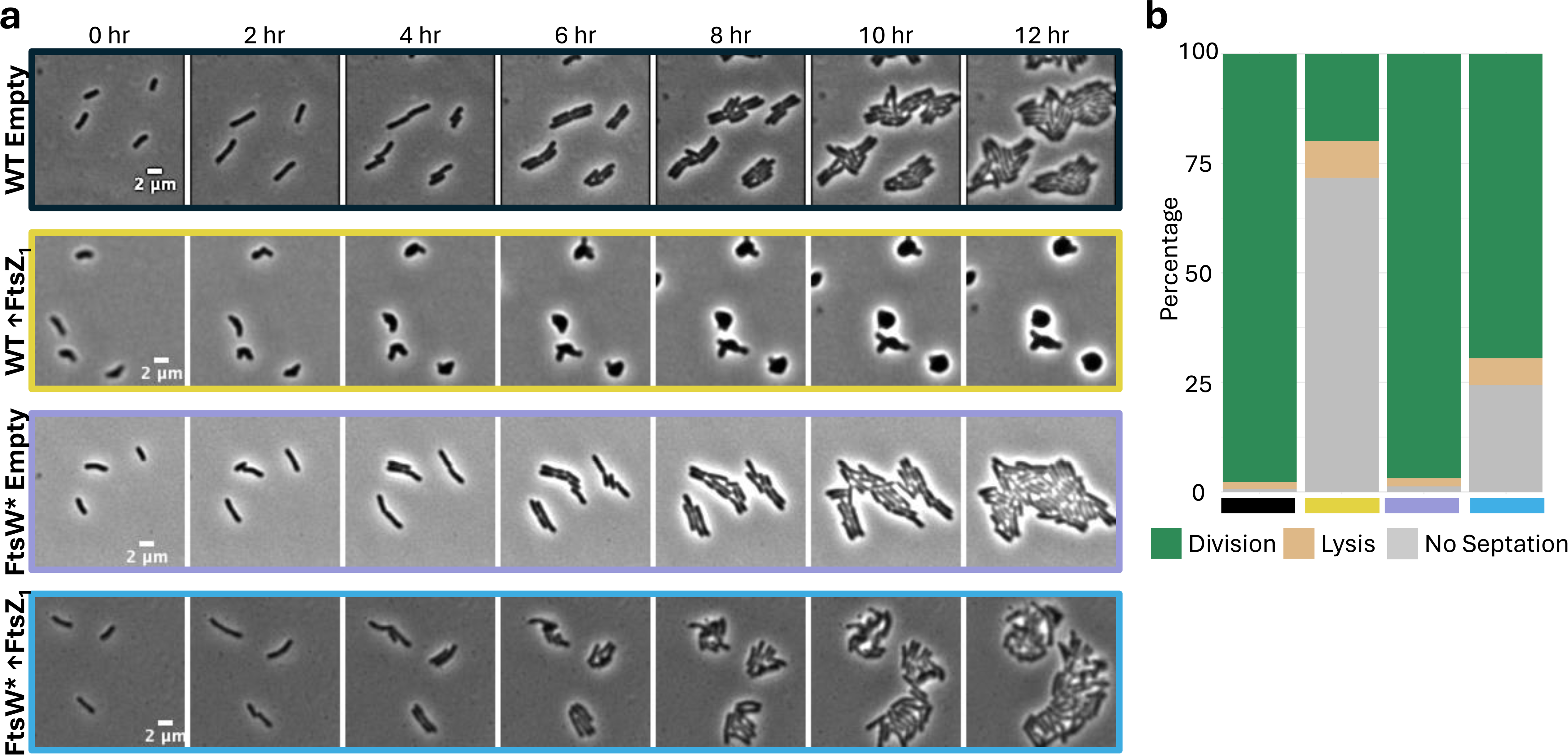
FtsZ_1_ inhibits the activation of septal peptidoglycan synthases which blocks septation. (a) Phase contrast microscopy of *A. tumefaciens* in the wildtype background with pSRKKM Ptac: empty vector and FtsZ_1_ as well as the FtsW* background with pSRKKM Ptac: Empty and FtsZ_1_. Images collected every two hours. (b) Characterization of mother cells from the images in the previous panel, at timepoint zero, for the categories of division, lysis, and no septation (n = 310, 311, 320, 308).

Interestingly, when grown on the agarose pads for this time-lapse experiment, the FtsW* background also partially rescued the cell morphology of cells overexpressing FtsZ_1_, with most cells retaining proper morphology with some mid-cell swelling. This discrepancy led us to speculate that decreased cell wall integrity could be responsible for the persistent morphological differences observed after 16 hours of FtsZ_1_ overexpression in the FtsW* background (Figure 5b). When grown in a liquid medium, bacteria encounter more direct and immediate osmotic challenges, which are known to change morphology and growth rates (Ye et al. 2018; Rojas et al. 2018). Thus, it is possible that osmotic differences exacerbate cell envelope mutant phenotypes when grown in liquid or on agarose or agar (Figure 5; Figure 6; Figueroa-Cuilan et al. 2021). Therefore, the same FtsW* strain experiences different stresses when induced for FtsZ_1_ overexpression for 16 hours in liquid medium verses on agarose pads. Overall, these data suggest that incorporation of excess FtsZ_1_ into the Z-ring likely misregulates septal peptidoglycan synthesis, leading to decreased cell wall integrity and susceptibility to osmotic pressure. These insights may hint at a physiological role for FtsZ_1_ as a tuning mechanism to ensure that the septal peptidoglycan is appropriately formed and remodeled to maintain the overall integrity of the cell wall during division. This may be particularly adaptive for microbes that reside within the rhizosphere or *in planta* where root exudates may shift the osmotic potential, requiring tuning of the cell wall structure.

Future work will be necessary to understand the molecular mechanisms underpinning the influence of FtsZ_1_ on cell division in the Hyphomicrobiales. Conservation of the GTP-binding pocket among duplicate GTPase domains within the Hyphomicrobiales order suggests that Z-ring incorporation is likely a shared feature (Figure 3cde). What remains to be determined is if any of the substitutions in the duplicate protein sequence are required for the observed FtsZ_1_ overexpression phenotype. Since *A. tumefaciens* belongs to the *Rhizobiaceae* family, it is possible that one or more of the colored regions in Figure 3cde are required for its regulatory function. If the shared yellow region is implicated, the *Bradyrhizobium* duplicate may also influence cell division as well. We hypothesize that these surface exposed regions may serve as interfaces for protein-protein interactions. Future efforts will focus on the identification of FtsZ duplicate binding partners. An improved understanding of the mechanisms and roles of the FtsZ duplicates in tuning cell division in plant-associated microbes could be exploited for biocontrol of agricultural pests or the enhancement of plant growth promoting microbes.

### How could duplicate FtsZ GTPases regulate septal peptidoglycan activation?

Two independent FtsZ duplication events produced secondary FtsZ GTPases that have been maintained primarily in groups with a plant-associated lifestyle within the Hyphomicrobiales order. While the deletion of FtsZ_1_ in *A. tumefaciens* was not detrimental in an array of conditions, overexpression led to misregulation of septal peptidoglycan biosynthesis. We hypothesize that FtsZ_1_, the *A. tumefaciens* ortholog of the duplicate FtsZ GTPases in the *Rhizobiacea* and *Phyllobacteracea* families, functions as a non-essential inhibitor of cell division which may be conditionally beneficial during microbe-plant interactions. Given the pattern of conservation, it is likely that it serves a similar role in regulating division in other family members.

The idea that Z-ring dynamics can regulate septal peptidoglycan synthesis is not a new concept. Across the bacterial domain, the Z-ring coordinates with the septal peptidoglycan machinery FtsWIQLB through Z-ring anchoring proteins such as FtsA (Park et al. 2021, Santiago-Collazo et al. 2024). In general, a subset of the divisome assembles with the Z-ring early in the process of division. This early divisome complex recruits late divisome components that will actively synthesize septal peptidoglycan, and correct assembly of all members serves as a checkpoint into late division. More specifically, FtsN has been shown to both interact with FtsA and activate the late divisome septal peptidoglycan synthases FtsW/FtsI (Addinall et al. 1997; Gerding et al. 2009; Park et al. 2021). A duplicate FtsZ could negatively impact septal synthase activation by directly or indirectly interfering with this checkpoint. The hyperactive FtsW* allele bypassed this effect because it can polymerize glycan strands without proper activation (Modell et al. 2014; Meeske et al. 2016; Rohs et al. 2018). Our observations that excess FtsZ_1_ inhibits division, and that FtsW* can alleviate this inhibition, suggest that FtsZ_1_ may disrupt late divisome activation.

FtsZ duplicates have already been shown to have regulatory roles in cyanobacteria and chloroplasts, where a FtsZ paralog functions to destabilize the Z-ring in a GTP/GDP- ratio-dependent manner (Porter et al. 2023; Porter et al. 2021). Recently, Cao *et al*. demonstrated that differences in the GTPase domains allowed the duplicates to reduce Z-ring stability in *Arabidopsis thaliana* chloroplasts (Cao et al. 2025). This work highlights how a duplicate FtsZ’s ability to increase the turnover of the Z-ring can impact septal synthase activation. As described by Hu et al., faster Z-ring assembly increases the likelihood of FtsN and FtsWIQLB interaction: therefore, activating septal peptidoglycan synthesis (Hu et al. 2025). Meanwhile, destabilizing the Z-ring would drive fewer interactions between FtsN and FtsWIQLB. This mechanism is different from what we propose for the FtsZ duplicates in the Hyphomicrobiales. Previous *in-vitro* work with *A. tumefaciens* proteins suggests that FtsZ_1_ does not affect Z-ring structure or turn- over (Howell et al. 2019). Instead, in this paper, we describe conserved amino acid sequences in the duplicate FtsZ GTPase domains that could scaffold protein-protein interactions. While further work is needed to discern if these sites drive or block specific interactions with the Z-ring, it is tempting to hypothesize that they function to disrupt late divisome recruitment, which is not yet well understood in the Hyphomicrobiales. Moreover, the putative FtsN ortholog, RgsS, in the *Phyllobacteriaceae/Rhizobiaceae* contains an extended cytoplasmic intrinsically disordered region and may have diverged in function and interacting partners from canonical FtsN (Krol et al. 2020). The differences in RgsS domain structure between the *Phyllobacteriaceae/Rhizobiaceae* and *Bradyrhizobiaceae* families (Krol et al. 2020) coupled with distinct regions of the duplicate FtsZ GTPase, may indicate that different binding partners activate FtsN/RgsS in the Hyphomicrobiales.

### Concluding Remarks

We are fascinated by the conservation of duplicate FtsZ GTPases in plant-associated bacteria. Plants often ward off potential infectious bacterial agents by secreting defensin-like peptides. Sublethal levels of these peptides have been shown to specifically block cell division and antagonize Z-ring function in *S. meliloti* (Li et al. 2002; Wu et al. 2012; Penterman et al. 2014; Heckel et al. 2014). Disruption of cell division triggers transcriptional changes via activation of the ChvGI two component system which has historically been associated with host invasion but have been more recently shown to provide protection against environmental stressors including antibiotics and antimicrobial peptides (Quintero-Yanes et al. 2022; Williams et al. 2022; Bouchier et al. 2025). It is intriguing to speculate that, at low levels, conditional expression of *ftsZ_1_* may inhibit or slow bacterial cell division: therefore, giving plant-associated bacteria time to counteract environmental stressors during host invasion, rhizosphere association, or other niche-specific interactions. Further research into these FtsZ GTPase duplications can help tease apart the complex relationship between expression, protein-protein interaction, and downstream function of these septal peptidoglycan inhibitors.

## Data Availability Statement

Strains and plasmids are available upon request. The authors affirm that all data necessary for confirming the conclusions of the article are present within the article, figures, and supplemental files.

## Acknowledgments

We thank all members of the Brown Laboratory for critical feedback on this article.

## Figure accessibility and alt text

Figure 1. A GTPase domain tree of Alphaproteobacterial FtsZ representatives shows that FtsZ duplicates are observed primarily in the Rhodospirillales and Hyphomicrobiales orders.

Figure 2. A cladogram of the families within the Hyphomicrobiales order is supplemented with counts of FtsZ duplicate-containing species for each family.

Figure 3. The GTPase domain of duplicate FtsZ sequences from the Hyphomicrobiales is represented with emphasis on conserved sequences found between the Nitrobacteraceae, Phyllobacteraceae, and Rhizobiaceae families.

Figure 4. Images of spotting assays and phase contrast microscopy show that FtsZ_1_ overexpression is toxic.

Figure 5. Images of spotting assays and phase contrast microscopy show that the FtsW* background resolves viability but not morphology of the FtsZ_1_ overexpression.

Figure 6. A series of phase microscopy images, two hours apart, show that FtsZ_1_ overexpression stops cell division in wildtype but not FtsW* cells.

